# Vocal mimicry in corvids

**DOI:** 10.1101/2025.03.26.645457

**Authors:** Claudia A.F. Wascher, Gemini Waterhouse, Bret A. Beheim

## Abstract

Vocal mimicry, the copying of sounds produced by another species or the environment, is commonly described in vocal learners, such as songbirds. Understanding the functions of vocal mimicry can help to uncover the evolutionary drivers of vocal learning. Different adaptive functions like interspecific and intraspecific communication have been suggested, as well as the possibility of vocal mimicry to be a mistake during vocal learning. In the present study, we review the occurrence of mimicry in the family of corvids and investigate the socio-ecological factors driving the evolution of vocal mimicry in this group of birds. We recorded evidence of vocal mimicry from primary (xeno-canto recordings) as well as secondary sources (published literature) and found evidence for vocal mimicry in 39 out of 128 corvid species (30%). Socio-ecological factors like breeding status, habitat and trophic niche did not have a significant effect. We used Bayesian modelling based on existing data from primary and secondary sources to estimate the occurrence of mimicry, suggesting that vocal mimicry may be more widespread among corvids than currently documented, with many species potentially being ‘hidden mimics’. Our study for the first time systematically reviews the occurrence of vocal mimicry across the family of corvids and investigates a range of socio-ecological factors driving the behaviour, hopefully inspiring future field work on those species.

## Introduction

Vocal mimicry, the copying of sounds produced by another species or the environment, is commonly described in songbirds (Kelley et al. 2008; Dalziell et al. 2015; Goller and Shizuka 2018) and also has been reported in mammals including an Asian elephant, *Elephas maximus* (Stoeger et al. 2012), a white whale, Del*phinapterus leucas* (Ridgway et al. 2012), a harbor seal, *Phoca vitulina* (Duengen et al. 2023), and greater mouse-eared bats, *Myotis myotis* (Ancillotto et al. 2022). Mimics usually acquire sounds through vocal production learning, the ability to produce novel signals through imitation or modification of existing vocalisations (Janik and Knörnschild 2021). Vocal production learning in animals has parallels with human vocal ontogeny (Ter Haar et al. 2021), as it requires an individual to process acoustic information from an appropriate model, engage in vocal practice and evaluation through auditory feedback, and requires enhanced forebrain control of the vocal organ (Nieder and Mooney 2020).

According to Dalziell et al. (2015), vocal mimicry is defined as production of vocalisations resembling heterospecific or environmental sounds which change the behaviour of the receiver after perceiving the acoustic resemblance between the mimic and the model, and the behavioural change confers a selective advantage on the species mimicking. This definition excludes other forms for vocal resemblance, for example as a result of chance, common ancestry (Goller and Shizuka 2018) or social learning (Bluff et al. 2010). Vocal mimicry may be a result of mistakes during vocal learning and have no function (Kelley et al., 2008). In contrast to this, a number of different potential functions of vocal mimicry have been previously suggested, broadly falling into two categories (a) interspecific communication, to avoid threats, dissuade competitors, or imitate threats to allow access to food resources and (b) intraspecific communication, in the context of sexual selection or for social affiliation (reviewed by Kelley et al., 2008). For example, mimics are suggested to produce vocalisations from predatory or agonistic species to deter competitors or potential predators from their territories (Dobkin 1979). Australian magpies, *Gymnorhina tibicen*, mimic several potential nest predators, for example the barking owl, *Ninox connivens*, and the boobook owl, *Ninox novaeseelandiae* (Kaplan 1999). Fork-tailed drongos, *Dicrurus adsimilis*, mimic alarm calls of other species to deter heterospecifics from food sources, in order to steal the food (Flower et al. 2014). Individuals might also benefit from vocal mimicry when used to attract other individuals. For example, greater racket-tailed drongos, *Dicrurus paradiseus*, mimic songs and contact calls to attract other species, which increases foraging efficiency (Goodale and Kotagama 2006a) and alarm calls of other species alongside their own mobbing calls, potentially attracting other species to support them when mobbing predators (Goodale and Kotagama 2006b). This shows that some species flexibly use vocal mimicry, and this may be context dependent. Finally, mimicked calls are sometimes incorporated into song or sexual displays and might play a role in sexual selection (Kelley et al., 2008). Overall, surprisingly little empirical research on the functional explanations of vocal mimicry is available.

Phylogenetic analysis has previously shown that vocal mimicry is unlikely to be an ancestral state in songbirds, meaning that it may have evolved independently on multiple occasions (Goller and Shizuka 2018). A review of European passerines estimated approximately 40% of songbird species to mimic (Garamszegi et al., 2007). Sounds mimicked by songbirds include birdsong (northern mockingbird, *Mimus polyglottos*: Gammon and Altizer 2011), alarm calls of other species (spotted bowerbird, *Ptilonorhynchus maculatus*: Kelley and Healy 2011; red-capped robin-chat, *Cossypha natalensis:* Oatley 1969; thick-billed euphonia, *Euphonia laniirostris*: Morton 1976), calls of birds of prey (blue jay, *Cyanocitta cristata*: Hailman 1990), and human voices (European starlings, *Sturnus vulgaris*: West et al. 1983). As suggested by Gammon (2013), the ‘model selection’, *i*.*e*. type of sound mimicked, can provide valuable insight into the potential functions of vocal mimicry. For example, northern mockingbirds, *Mimus polyglottos*, preferentially mimic species whose sounds are acoustically similar to their non-imitative song, supporting the acoustic similarity hypothesis, and suggesting a non-adaptive explanation of vocal mimicry (Gammon, 2013). Mimicry of predator calls by Steller’s jays, *Cyanocitta stelleri*, is also linked to mate selection, with mimicry occurring more frequently during the breeding season and when in the presence of their mates (Tippin 2017).

Corvids *(Aves: Corvidae)*, are a large and diverse group with over 120 species split between 21 genera, an array of which can be found on all continents except Antarctica (Madge and Burn 1994). Many corvid species have demonstrated advanced cognitive abilities in the social and physical domain (Taylor 2014). Corvids belong to the suborder of oscine passerine birds, where the vocal organ (syrinx) is highly developed to produce vocalisations (Suthers and Zollinger 2004). Vocalisations in corvids present a wide range of ecological and social adaptations, such as communicating information about predators such as predator type or level of risk or individual recognition (Wascher and Reynolds 2025). Corvids are open-ended vocal learners, which means they can acquire new vocalisations throughout their lives and not only in a specific sensitive period (Brenowitz et al. 1997). Carrion crows are capable of volitional control of vocalisations, *i*.*e*. can be trained to vocalise and withhold vocalisations in response to arbitrary stimuli (Brecht et al. 2019). Vocalisations in New Caledonian crows, *Corvus moneduloides* (Bluff et al. 2010) and common ravens, *Corvus corax* (Enggist-Dueblin and Pfister 2002) are socially learned from conspecifics. Several species of corvids have demonstrated the ability to mimic human speech when hand-raised and kept in close proximity to humans (Bluff et al., 2010). Mimicry of predator calls has been documePhylogenetic analysis has previously shown that vocal mimicrynted in several corvid species including Steller’s jays (Hope 1980; Tippin 2017), Sri Lanka blue-magpies, *Urocissa ornata* (Ratnayake et al. 2010), Florida scrub-Jays, *Aphelocoma coerulescens* (Woolfenden and Fitzpatrick 2020), blue jays, *Cyanocitta cristata* (Hailman 1990), and Eurasian jays, *Garrulus glandarius* (Goodwin 1951). Tippin (2017) conducted an in-depth study of vocal mimicry in Steller’s jays, which found significant differences in age and sex between mimicking and non-mimicking individuals, where males were less likely to mimic compared to females, and the likelihood of an individual mimicking decreased with age.

Missing data present a significant challenge in the study of animal behaviour, limiting our understanding of key phenomena (Garamszegi and Møller 2012; Nakagawa 2015). Factors such as habitat inaccessibility (Evans et al. 2018), species rarity (Roberts et al. 2016), human-centric biases in study design (Archer et al. 2014; Ellison et al. 2021; Bowler et al. 2025; Daw et al. 2025), and technological constraints (Hebblewhite and Haydon 2010) contribute to missingness. In evolutionary biology, missing data can skew our understanding of trait evolution, obscuring the selective pressures and socio-ecological contexts that drive behavioural adaptations. The problem of missing data can be addressed with more systematic data collection, e.g., planned missing data design (Noble and Nakagawa 2018), application of emerging technologies and statistical approaches, such as imputation procedures (Augustine et al. 2014), and a recognition that the absence of evidence is not evidence of absence in behavioural research.

The aim of the present study is to describe the occurrence of mimicry in corvids. We investigated the evidence for mimicry from primary, i.e. recordings on xeno-canto, and secondary sources, i.e. published literature. Further, we investigate socio-ecological factors affecting the occurrence of vocal mimicry in corvids. Vocal mimicry in corvids might serve a function of interspecific communication, such as deterring a potential predator and competitors, by copying the vocalisations of animals that are predatory or agonistic (Dobkin 1979). If this is the case, we expect smaller corvid species to mimic more, as they are more at risk of predation (Dierschke 2003) and we expect them to mainly mimic birds of prey. If vocal mimicry serves intraspecific communication, such as sexual selection (Kelley et al. 2008; Dalziell et al. 2015), we expect a positive correlation between the size of the vocal repertoire and the occurrence of vocal mimicry. Additionally, we investigate whether habitat, trophic niche and breeding system affect the occurrence of mimicry in corvids, although we do not have specific predictions in this regard and consider our analysis exploratory. In addition to biological factors driving vocal mimicry in corvids, species differ significantly in terms of study effort (Wascher & Reynolds, 2025). We investigate how differences in recording effort affect the detection of mimicry in corvids and evaluate the likelihood of ‘hidden’ mimicry.

## Methods

### Data on vocal mimicry

The present study was approved by the School of Life Sciences departmental ethics panel at Anglia Ruskin University (ETH2223-3747). This is a desk-based study, and no animals have been used for the research. Data were compiled from a variety of primary (xeno-canto) and secondary sources (peer-reviewed journal articles and books). Scientific and common names were standardized using a combination of the IOC Bird list version 14.1. (Gill et al. 2024) and the ‘Clements checklist’ (Clements et al. 2024) and 128 species of corvids were included. A xeno-canto advanced search was conducted between the 27^th^ of April and the 1^st^ of May 2023 (searches have been updated on the 22^nd^ of June 2023 and 5^th^ May 2024). Xeno-canto is a citizen science project and repository in which volunteers record, upload and annotate recordings of bird calls (www.xeno-canto.org, last accessed on 29^th^ July 2025). The xeno-canto search yielded a total of 19,566 audio recordings of corvid calls from 128 different species (mean±SD per species: 153 ± 360), belonging to 21 genera (mean±SD per genus: 889 ± 1,767). The number of recordings per species ranged from zero to 2,272 recordings. All recordings with mentions of ‘mimicry’, ‘imitation’ or ‘production of heterospecific calls’ in the comments or labels section were identified. For the purpose of this study, we defined vocal mimicry as the occurrence of any type of non-conspecific sound and a resemblance between the mimicked and model sound (Kelley et al. 2008; Goller and Shizuka 2018). Our definition does not require the receivers to respond behaviourally to mimicked sound in a way that benefits them, which is the case for the definition of vocal mimicry in Dalziell et al. (2015). We noted the duration of the recording, date, latitude and longitude and type of sound mimicked. Recordings were then listened to by GW and CAFW, and if both observers agreed that they contained convincing evidence for production of mimicry the observation was considered as evidence of mimicry. We fully acknowledge that this process involves a high level of subjectivity and a risk of false positives, i.e., the recordings not being a case of mimicry in corvids. In order to account for this, we have rated the perceived ‘reliability’ of the recordings, indicating whether credibility of the recording was high, moderate or low. A total of 466 records of mimicry have been identified. For 350 records it was indicated that the vocalising bird was seen by the recorder, for 86 records the bird was not observed and for 30 records this information was unknown. The majority of records (325) have been classed as ‘high’ reliability, meaning that the rater (CAFW) had high confidence in the recording containing mimicry by a corvid, 53 records were classed as ‘moderate’ reliability and 88 records as ‘low’ reliability, meaning the record is more likely to be a false positive.

Data were compiled from a range of primary and secondary literature sources. A search was conducted using Google Scholar, combining search terms such as ‘mimic’, ‘imitate’, and ‘copy’ with both English and Latin species names. In addition, we consulted authoritative field guides and handbooks, including for example the *Handbook of the Birds of the World* and *Birds of the World*, which were manually reviewed for references to vocal mimicry or imitation. Only sources that explicitly used terms such as “mimic” or “imitate” and clearly identified the mimicking species and the target of mimicry (e.g., species, family, order, or group such as ‘environmental sounds’) were included. We excluded any instances where vague descriptors like ‘similar to’ or ‘reminiscent of’ were used. We have included reports of mimicry from six books and field-guides (non-peer reviewed sources) and 11 peer-reviewed journal articles. A full list of sources is provided in the online supplementary material.

For each species, the number of entries in the corvids literature database (Droege and Töpfer 2016) plus one entry in Birds of the World (Poole & Gill, 2020) was used as an estimate of the volume of secondary literature available (range: 1-423 entries). For each piece of evidence of vocal mimicry, CAFW allocated a ‘reliability score’ of high, moderate or low. Sixty-five percent of recordings have been rated as ‘high’, 23% as ‘moderate’ and 12% as ‘low’ reliability.

For each of the 128 species, we recorded body mass (information available for 121 species), vocal repertoire (98 species), breeding system (cooperative breeding, groups, territorial pairs; reported for 121 species), habitat (coastal, desert, forest, grassland, human modified, rock, shrubland, woodland; reported for 124 species) and trophic niche (frugivore, granivore, invertivore, omnivore; reported for 124 species).

### Statistical analysis

All analyses were conducted in R version 4.3.1. (R Core Team 2019). We constructed a phylogenetic tree for the Corvidae family using data from the Open Tree of Life (OTL). The tree was retrieved from the Open Tree of Life database (OpenTreeOfLife et al. 2019) using the rotl R package (Michonneau et al. 2016). We employed Bayesian generalized linear mixed-effects models built with the MCMCglmm package (Hadfield 2010) to determine effects of body size, vocal repertoire size, habitat and trophic niche on vocal mimicry in corvids. Genus was included as a random factor and the OTL tree served as the foundational phylogeny for the MCMCglmm. We ran Markov chain Monte Carlo (MCMC) chains for 3,000,000 iterations, thinned by 2000, and employed a burn-in of 2,000,000. All main effects were given normal(0, 0.5) log-odds prior, with a normal(0, 1) log-odds intercept prior. Random effects by genus were given an inverse Wishart prior with V = 1 and nu = 10. Data and code of the present study are published as online supplementary materials (Supplementary File 2; supplementary File 3).

In order to estimate the occurrence of mimicry, we have developed a Bayesian model of evidence updating, assuming that for any specific kind of evidence (citation database entries, audio recordings), there is a corresponding rate at which mimics are detected per-unit evidence. The model enables the calculation of the posterior probability a species in which no mimicry is detected is, in fact, a mimic, which is a more principled way to handle the presence of zeros in such datasets than taking them at face value (Beheim et al. 2021; Sigmundson et al. 2025). Full details of the model are presented in the supplementary materials, including several robustness checks for variable reporting rate and quality across species, insensitivity to priors, and the effects of reliability scores on evidence accumulation. Results of these checks are presented in the online supplementary materials. The present manuscript was posted as pre-print and full statistical analysis code is available in the accompanying materials repository (Wascher et al. 2025; https://github.com/babeheim/corvid-mimicry).

## Results

### Description of the dataset

We recorded evidence of vocal mimicry in 39 species of *Corvidae*, belonging to thirteen genera (30% of species; Figure 1). Thirty-three species of corvids are described as mimics in the literature and recordings from 17 species have been identified on xeno-canto, including six species not previously described as mimics (Indochinese green-magpie, *Cissa hypoleuca;* carrion crow, *Corvus corone*; black-chested jay, *Cyanocorax affinis*; Yucatan jay, *Cyanocorax yucatanicus;* clark’s nutcracker, *Nucifraga Columbiana*; rook, *Corvus frugilegus*; Table 1). One to 71 different types of sounds were mimicked (mean ± standard deviation: 5.512 ± 11.568). Sounds have been mimicked from a wide range or birds (Accipitriformes, Anseriformes, Charadriiformes, Cuculiformes, Falconiformes, Galliformes, Gaviiformes, Gruiformes, Pelecaniformes, Passeriformes, Piciformes, Strigiformes), mammals (Artiodactyla, Carnivora, Perissodactyla, Primates (human voice), Rodentia) amphibian (Anura) and insect species (Hemiptera) as well as anthropogenic sounds, such as bells, lawnmower, motorbike (see supplementary materials for full list of mimicked species). Body mass, repertoire size, habitat, breeding system and trophic niche did not affect mimicry status (Table 2).

**Table 1:** Cross-table indicating the number of species detected as mimics in the literature and on xeno-canto. Records per species indicate the number of records in the corvid literature database and xeno-canto.

| mimicry category | n.species | number of records in the corvid literature database | number of records on xeno-canto |
| --- | --- | --- | --- |
| no mimicry seen | 89 | 11 | 51 |
| number of species detected as mimics in the literature only | 22 | 29 | 203 |
| number of species detected as mimics on xeno-canto only | 6 | 126 | 477 |
| number of species detected as mimics both in the literature and on xeno-canto | 11 | 94 | 697 |

**Table 2.**
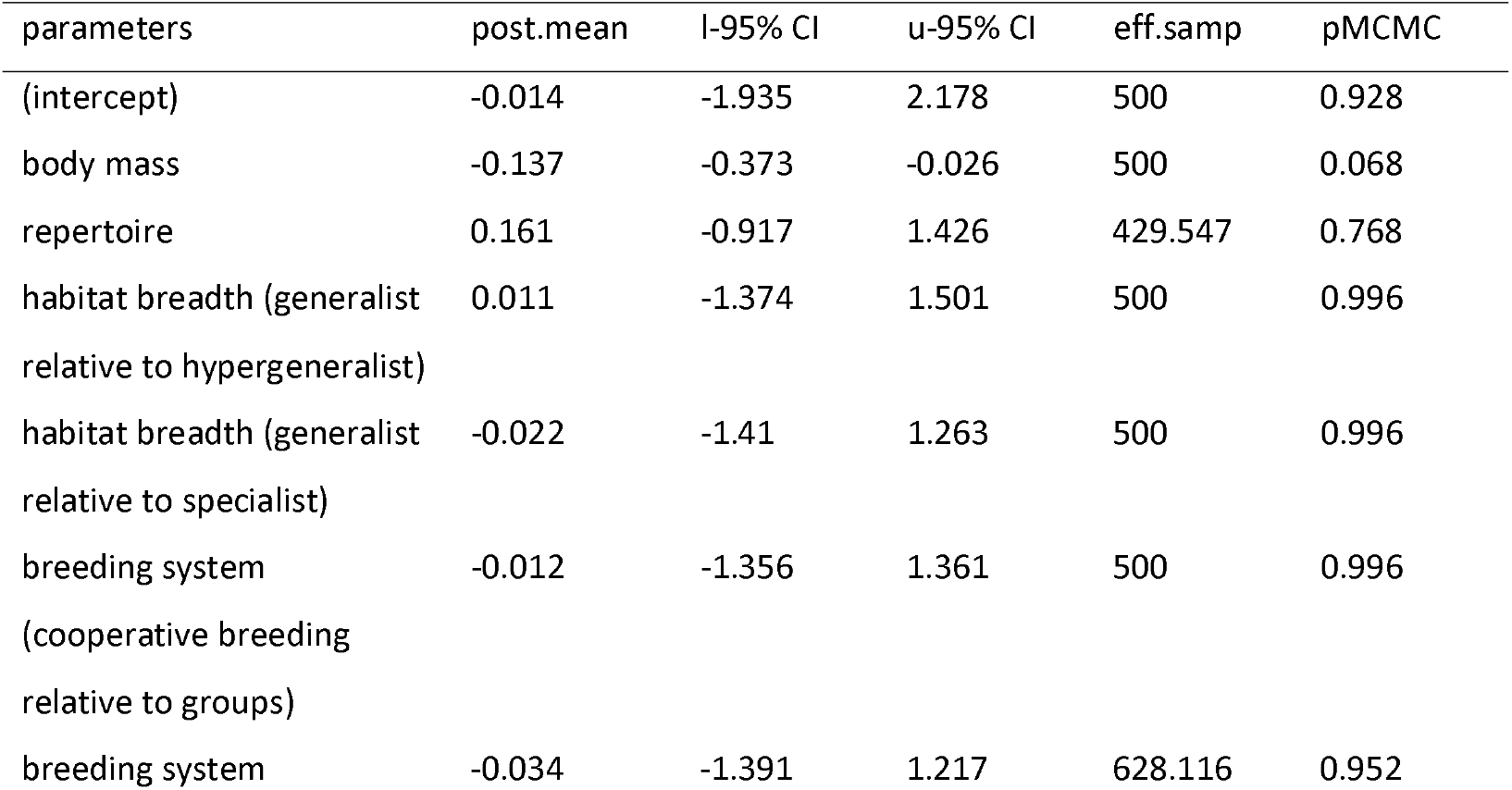
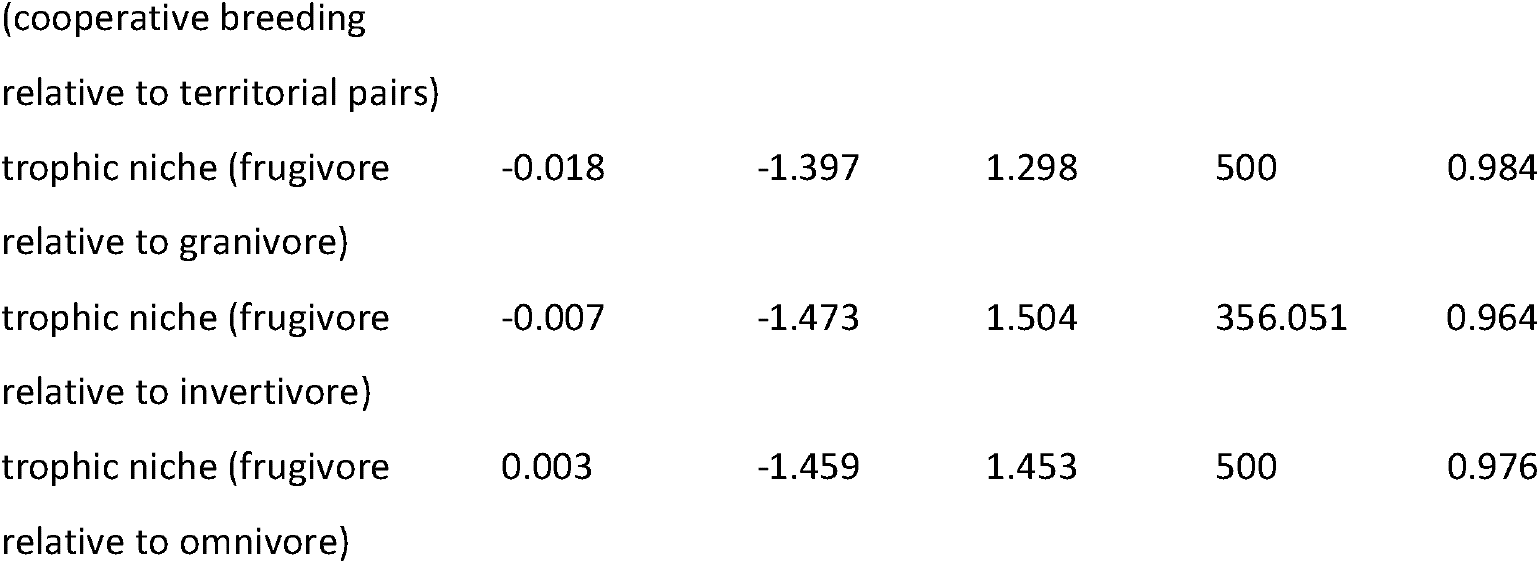
Results of Markov chain Monte Carlo generalized linear mixed-effect model (MCMCglmm). Factors with significant effects (*p* ≤0.05) are shown in bold.

**Figure 1.**
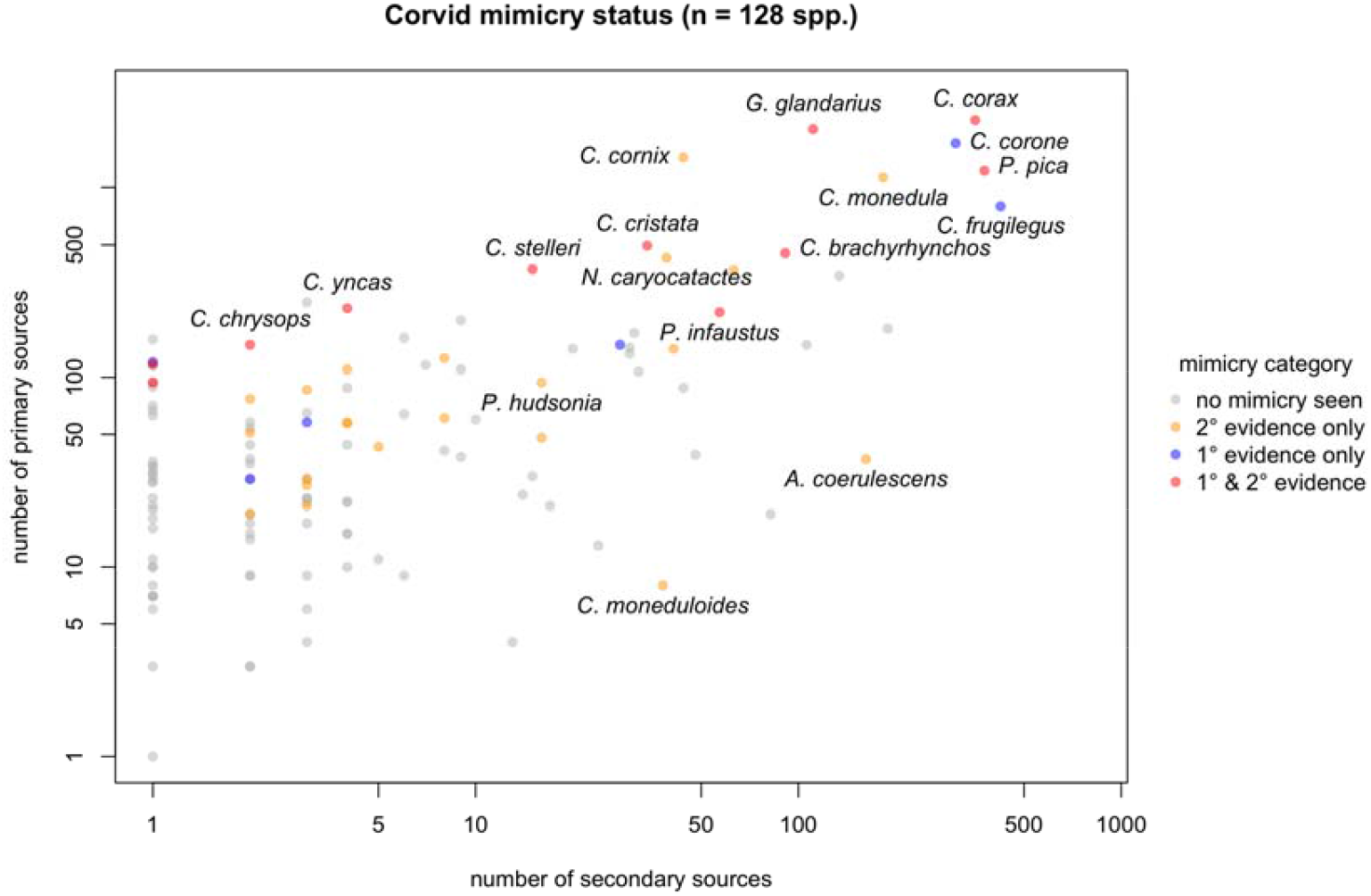
Vocal mimicry in 128 species of corvids. Grey points indicate species with no mimicryin the literature, blue points illustrate species with recordings of mimicry on xeno-canto but no records in the literature and yellow dots indicate species only reported as mimics in the literature, but no recordings of mimicry are available on xeno-canto.

### Estimating the occurrence of mimicry

Because species with more evidence are more likely to be reported as mimics, the observed rate of occurrence of mimicry in the published literature (33/128 species) and in the xeno-Canto recordings (17/128 species) are likely to undercount the true occurrence of mimicry, as species that are capable of mimicry have not (yet) been documented as such. In order to formalize this intuition, we developed a Bayesian model of evidence accumulation that can accurately estimate the true rate of occurrence of a phenomenon with variable rates of reporting, detection and source quality (see the supplementary information for full specification). Figure 2 shows how different information sources update prior assumptions about the occurrence of mimicry across corvids. When both primary and secondary sources are incorporated into the model, each with their own detection rates, we estimate that mimicry occurs in 82% (+/-7%) of *Corvidae*. Using this model, we can also calculate a composite evidence score, in practice converting a count of recordings on xeno-canto into the equivalent number of corvid database entries (or vice versa), which is useful to understand how the hidden mimicry probabilities for each species relates to research attention (Figure 3). When very little is known about a particular species (e.g. no uploaded recordings or entries in the citation database), our model assumes that the best guess about their mimicry ability is the corvid-wide intercept in the model, q. As information accumulates with no positive detection of mimicry, the species-specific posterior estimate declines according to the mathematical relationship described in the supplement. Thus, although most species are not detected as mimics, only a handful actually have sufficient evidence to update species-specific posteriors near zero probability (Figure 3).

**Figure 2.**
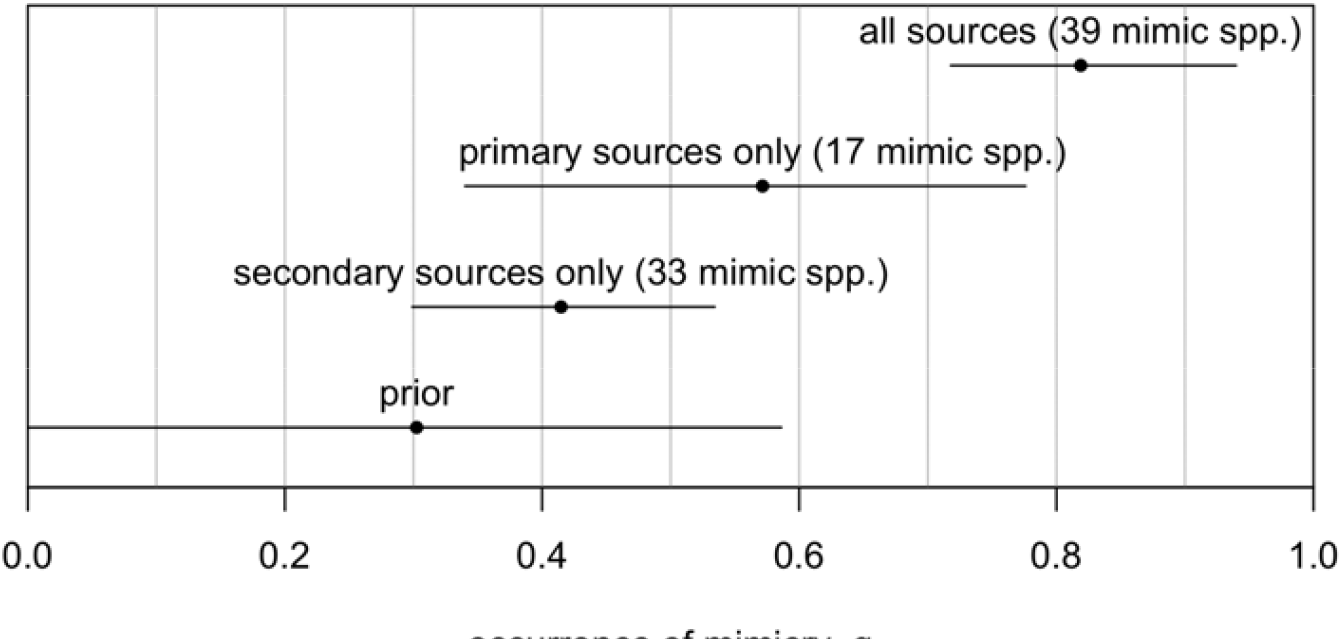
Prior and posterior estimates (means and 89% HPDI) of the frequency of occurrence of mimicry across corvid species conditioning on different information evidence (primary sources, secondary sources, and both combined).

**Figure 3.**
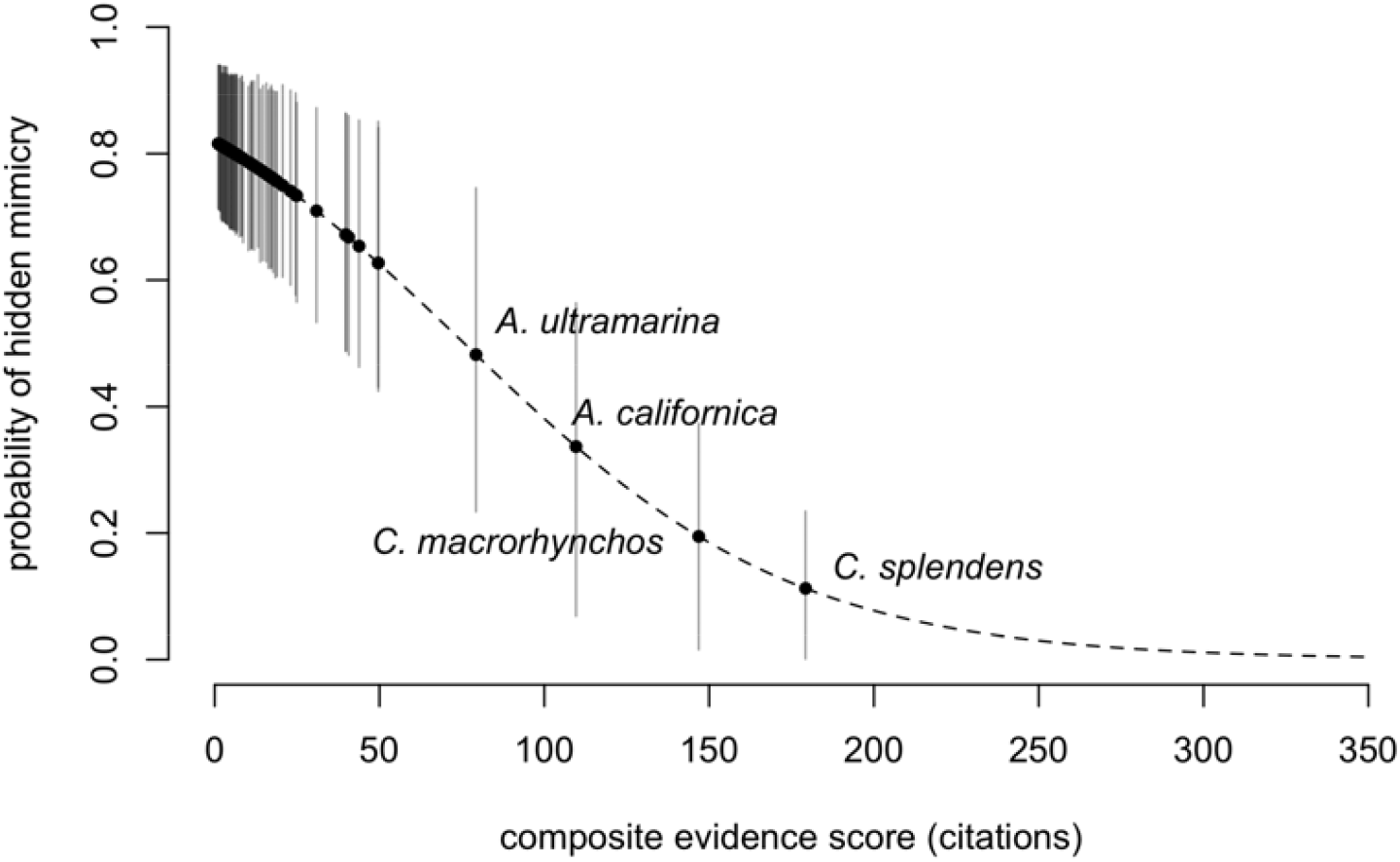
Composite evidence score related to the probability of hidden mimicry.

Looking at posterior probabilities of hidden mimicry for each species individually. The ten species which are predicted as least likely to be mimics and ten species nominated as most likely to be mimic are presented in the supplementary materials.

## Discussion

In the present study, we describe cases of vocal mimicry in 39 species of corvids belonging to thirteen genera, showing that vocal mimicry is widespread across this family of birds. With 30% of corvids known to mimic, it is more prevalent in this group, as opposed to the 8.9% observed in the entire suborder of songbirds (Goller and Shizuka 2018). Additionally, because mimicry detection is strongly associated with the amount of research effort directed at any specific corvid species, there is reason to believe that the occurrence of vocal mimicry is starkly underestimated by the current empirical evidence and many species of corvids are likely to be ‘hidden mimics’, meaning they are undetected as mimics by the currently available literature and data.

Our study underscores the importance of crowd-sourced databases, such as xeno-canto, revealing potentially overlooked or underreported behaviours, such as vocal mimicry. We have identified five species of corvids as mimics based on xeno-canto recordings, which have not previously been reported as mimics. Xeno-canto has previously been used scientifically to develop machine learning methods for automatic detection and classification (Ovaskainen et al. 2018), describing geographic variation (Deng et al. 2019), and interspecific variation (Benedetti et al. 2018). Our analysis also highlighted the utility of combining primary and secondary data sources in assessing the occurrence of mimicry. By using a Bayesian model to integrate these data, we were able to generate more probabilistic estimates of mimicry occurrence across species, reducing uncertainty in our findings. The composite evidence score, which converts xeno-canto recordings into a standardized measure comparable to traditional literature-based records, provides a valuable tool for assessing the hidden probabilities of mimicry within under-studied species. This model also offers insights into how research attention and available data influence our understanding of mimicry, suggesting that species with more extensive documentation are more likely to be perceived as mimics, while those with less attention may still exhibit mimicry but remain underreported. These calculations are useful starting places for future fieldwork focusing on vocal mimicry and can already be subjected to an empirical test.

Another novel aspect the Bayesian modelling allowed us to do is performing several robustness checks to our interpretation of data. For example, supplying the empirical estimates and evidence distribution as a simulation input for 1,280 corvid species, which presents ten times the real number of species in the group, the posterior estimates of occurrence of mimicry consistently approach the true value of each data sources (primary only, secondary only, primary and secondary). Potentially low data quality, is a persisting challenge in citizen science projects (Kosmala et al. 2016), including the use of data from xeno-canto. For example, misclassification and ‘noisy’ data can significantly hinder application of machine learning on citizen science datasets (Gupta and Gupta 2019). However, depending on type of application, unscreened crowd contributors also can be as effective as experts in data annotation (Cocos et al. 2017). In order to evaluate the impact of potentially low data quality, potentially increase the number of false positives and hence us over-estimating the occurrence of mimicry in corvids, we have scored the perceived reliability of each report of vocal mimicry as high, moderate or low. Most recordings were rated as high reliability; however, in three species only low reliability recordings were available, hence increasing the likelihood of type I error. To check for sensitivity of false positives, we built models under the assumption that all low reliability records are false and another model that only high reliability records are true. Changes in the model did not substantially shift the estimate of occurrence, which remained above 80% in the most conservative false positive models. This means we can assume the evidence presented in the current study to be robust.

We did not find body mass or other socio-ecological factors such as breeding system or habitat to affect mimicry status, hence we cannot draw conclusions about potential functions of vocal mimicry in corvids. Anecdotal reports in the literature suggest a potential ‘deterrent’ function, e.g. anti-predator or to decrease competition. For example Steller’s jays mimicked hawk calls while being mobbed by hummingbirds, as reported in the notes on xeno-canto. Similarly, green jays, *Cyanocorax yncas*, were reported to disperse a foraging group of plain chachalacas, *Ortalis vetula*, whereupon jays moved in to consume seeds being eaten by *chachalaca* (Gayou 2020). In Canada jays most sounds mimicked are known or potential predators of adults or nests, hence mimicking predators is assumed to serve as a warning call to other group members, to confuse the predator itself, or merely to signal in a general way that a threat is present (Strickland and Ouellet 2020). Blue jays, *Cyanocitta cristata*, have been observed to mimic several species of birds of prey and other bird species, in order to make other species to leave food sources, *e*.*g*., feeder and therefore reduce competition (Hailman 2009). This suggests vocal mimicry in corvids to have an adaptive evolutionary function and heterospecific sounds might not, as previously suggested, be learned by mistake (Kelley et al. 2008). However, in order to investigate functional contexts of vocal mimicry behavioural observations of context or experimental tests on behavioural responses of receivers are necessary. An overabundance of mimicry of sounds from alarm and aggressive contexts has been also described for example in spotted bowerbirds, *Ptilonorhynchus maculatus* (Kelley and Healy 2011). Fork-tailed drongos have been described to mimic in order to deceive heterospecifics, for example to lead pied babblers, *Turdoides bicolor*, and meerkats, *Suricata suricatta*, away from food sources in order to steal the food (Flower et al. 2014). Mimicry by female birds has been observed to deter nest predators and protect eggs and nestlings (Igic et al., 2015).

Corvids mimicked a wide range of different sounds, including calls from birds of prey or other non-predatory bird species, human speech, and mammalian vocalisations. The fact that corvids are capable of mimicking such a wide range of sounds further highlights their cognitive flexibility and auditory learning abilities. Interestingly, in a number of species corvids have been reported to mimic human speech in captivity (Landsborough Thomson 1964; Lorenz 2002), whereas such reports are rare from wild birds. This further supports the high level of flexibility in mimicry. We suggest that while many, if not all corvid species possess a general capacity for vocal mimicry, the extent to which this ability is expressed in the wild likely depends on socio-ecological factors such as predation pressure and competition.

Our study was of explorative nature, our capacity to test specific hypotheses regarding the evolutionary drivers of mimicry in corvids was limited. Vocal mimicry has previously been described to be driven by social factors. Parrots for example preferentially mimic socially relevant vocalisations (Bradbury and Balsby 2016). In our study, we showed that the breeding system (territoriality, family groups and cooperative breeders) does not seem to affect the occurrence of mimicry. This does not mean that social factors do not affect vocal mimicry in general. Corvids generally are a group of highly social birds (Wascher, 2018) forming strong social relationships within their groups (Fraser and Bugnyar 2010) and memorizing vocalisations of social partners for several years (Boeckle and Bugnyar 2012). Because of its flexibility, the social system of corvids is difficult to characterize, for example a number of species are facultative cooperative breeders, where the occurrence of cooperative breeding depends on environmental factors. Additionally, the social system depends on life-history factors, for example individuals form large flocks outside of the breeding season or when they are young and become territorial, when they mate and start to reproduce (Wascher 2018). We suggest that because we did not find an effect of the breeding system on the occurrence of vocal mimicry, this might indicate sexual selection not to be a main evolutionary driver of vocal mimicry in corvids. Sexual selection has been proposed as one of the potential selective drivers of vocal mimicry, especially in birdsong, where mimicked vocalisations could provide an honest indicator of male quality (Dalziell et al. 2015).

To summarize, our study contributed to a broader understanding of vocal mimicry. Vocal mimicry in birds has been described to have evolved several times in different families and independently of vocal learning (Goller and Shizuka 2018). A better understanding of the socio-ecological drivers of the behaviour offers valuable insights into the evolution of communication systems in birds and other animals. By investigating mimicry in corvids, we gain an opportunity to explore how complex vocal behaviours evolve, including vocal learning.

## Acknowledgement

We thank all xeno-canto members for sharing their recordings and two anonymous referees for their constructive advice and helpful comments on our manuscript.

