## Supplementary materials for "Vocal mimicry in corvids"

### Estimating the Occurrence of Mimicry

#### A Bayesian model of knowledge

To begin, we assume that for any specific count of evidence, there is a rate $\rho$ at which mimics are detected per-unit evidence.

Conceptually, this is an *average* detection rate across mimic species and recordings, rather than the rate for a specific species. It is a property of (a) the rate of display of mimicry behavior for the sample of birds under observation and (b) also the accuracy of detection by people tagging the calls.

A critical assumption is that *the amount of evidence is independent of mimicry status*. That is, people are not focusing more on birds because they are mimics. Substantial taxonomic biases do exist, which correspond to clear geographic biases for species physically accessible to Western academic researchers. However, the causal justifcation is clear.

Having made this assumption we can treat the model as a Bayesian updating process in which the probability a particular species is a mimic conditional on the available evidence $k_{i}$ and detection rate $\rho$ is something we can estimate from data using Bayes’ Rule (Sigmundson, et al. *2025*).

Let $q$ be the true frequency of mimicry across corvids, such that any specific corvid is a mimic with probability $q$. Within the model, this serves as a Bayesian prior on our state of knowledge. Let $M_{i}$ be the event that species $i$ is really a mimic, and $D_{i}$ the event that mimic behavior was detected. Then it follows that a mimic species will not be detected after $k_{i}$ recordings with probability $\left( 1-\rho\right)^{k_{i}}$, and so will be a “hidden mimic”. Conversely, the probability of being detected after $k_{i}$ recordings is $1-\left( 1-\rho\right)^{k_{i}}$. It follows that:

$$Pr\left( D_{i}|M_{i} \right)=1-\left( 1-\rho\right)^{k_{i}}$$

$$Pr\left( D_{i}|M_{i}^{c} \right)=0$$

$$Pr\left( M_{i} \right)=q$$

Hence, by Bayes’ theorem,

$$Pr\left( M_{i}|D_{i} \right)=1$$

More interesting to us is the probability when detection does *not* occur:

$$Pr\left( M_{i}|D_{i}^{c} \right)=\frac{\left( 1-\rho\right)^{k_{i}}q}{\left( 1-\rho\right)^{k_{i}}q+\left( 1-q \right)}$$

This describes the probability a species in which no mimicry is detected is, in fact, a mimic, which we call the rate of *hidden mimicry*.

#### Applying this to empirical data

Since $k_{i}$ and $D_{i}$ are known, we can fit the remaining parameters of the above model in a Bayesian framework by assigning Beta priors to $\rho$ and $q$:

$$\rho\sim Beta\left( 1,3 \right)$$

$$q\sim Beta\left( 2,2 \right)$$

We then update the model using the observed data for all 128 species simultaneously to calculate posteriors by Hamiltonian MCMC (via R v4.4.2 using the {cmdstanr} library v0.8.1), sampling each model for 8,000 iterations over 4 chains. Convergence was confirmed for all models by Rhat < 1.01 and visual inspection of chains. Outcomes of models are presented in Table S1 and S2.

Table S1: Posterior Stan estimates using the single-source model using data from primary sources (xeno-canto recordings).

| parameter | mean | sd | lb | ub |
| --- | --- | --- | --- | --- |
| lp__ | -43.66 | 1.01 | -44.89 | -42.64 |
| q | 0.5715 | 0.1390 | 0.3401 | 0.7760 |
| rho | 0.0037 | 0.0019 | 0.0012 | 0.0059 |

Table S2: Posterior Stan estimates using the single-source model with secondary sources (Corvid database counts).

| parameter | mean | sd | lb | ub |
| --- | --- | --- | --- | --- |
| lp__ | -72.55 | 0.98 | -73.76 | -71.54 |
| q | 0.4148 | 0.0748 | 0.2990 | 0.5343 |
| rho | 0.2528 | 0.0973 | 0.1116 | 0.4013 |

Comparing estimates of rho for each information source shows the secondary literature indicates a much higher detection rate per unit evidence (database entries) compared to the primary sources (audio recordings). The q parameter in the secondary lit estimates a lower baseline rate than the primary lit, though *both* occurrence rates are much higher than the observed *Corvidae* mimicry frequency of 30% (39/128).

Given these posterior estimates for the model parameters, we can also calculate a *posterior probability of being a hidden mimic* for each species who was not detected as a mimic, $Pr\left( M_{i}|D_{i}^{c} \right)$, using Bayes’ rule. Before doing that, however, it would be useful to consider how we can use *both* information sources, primary and secondary, within the same modeling framework, which is a task we turn to next.

#### Expanding the model to include two evidence streams

One way to conceptualize a composite task is to first update the probability of mimicry on one evidence source ($k_{2}$) and then on a second evidence source ($k_{1}$), treating the output of one calculation as the input of the next. As above, if *either* evidence source detects mimicry, we assume that the probability of being a mimic goes to 1, so

$$Pr\left( M_{i}|D_{1i}\cup D_{2i} \right)=1$$

The probability a species $i$ is a hidden mimic after conditioning to both information sources ($k_{1i}$ and $k_{2i}$), conversely, can be expressed as

$$Pr\left( M_{i}|D_{1i}^{C}\cap D_{2i}^{C} \right)=\frac{q\left( 1-\rho_{2} \right)^{k_{2}i}\left( 1-\rho_{1} \right)^{k_{1}i}}{q\left( 1-\rho_{2} \right)^{k_{2}i}\left( 1-\rho_{1} \right)^{k_{1}i}+\left( 1-q \right)}$$

As before, we can specify Beta priors on $\rho_{1}$, $\rho_{2}$ and $q$ and update this model using Stan. Fitting on the full dataset of 128 species produces the following table of estimates (Table S3).

Table S3: Posterior Stan estimates using the two-source model with both primary and secondary sources included.

| parameter | mean | sd | lb | ub |
| --- | --- | --- | --- | --- |
| lp__ | -141.95 | 1.22 | -143.54 | -140.43 |
| q | 0.8190 | 0.0703 | 0.7178 | 0.9397 |
| rho1 | 0.0017 | 0.0004 | 0.0010 | 0.0023 |
| rho2 | 0.0197 | 0.0041 | 0.0134 | 0.0261 |

As before, the primary lit rho1 is much lower than the secondary lit rho2. The estimate of the Corvid-wide rate of occurrence, $q$ is now exceptionally high, implying that nearly 3/4ths of all species are expected to be mimics.

We can see how the combination of information sources updates $q$ in main text Figure 2.

### Interpretation of results

We can use the results of the above model in three ways: 1. calculating the species-specific probability of hidden mimicry 2. imputing datasets of mimicry for ancestral state reconstruction 3. incorporating the evidence for mimicry inside regression models of its global occurrence

#### A composite evidence score

In the two-channel model, we can express the amount of evidence from one channel in terms of the amount of evidence from another channel, the rho parameters, the prior, and a given posterior probability. Generically:

$$q_{i}'=\frac{q\left( 1-\rho_{1} \right)^{k_{1i}}\left( 1-\rho_{2} \right)^{k_{2i}}}{q\left( 1-\rho_{1} \right)^{k_{1i}}\left( 1-\rho_{2} \right)^{k_{2i}}+\left( 1-q \right)}$$

where $q_{i}'=Pr\left( M_{i}|D_{1i}^{c}\cap D_{2i}^{c} \right)$. Rearranging gives

$$k_{2i}={log}^{-1}\left( 1-\rho_{2} \right)\left[ log\left( \frac{q_{i}'\left( 1-q \right)}{\left( 1-q_{i}' \right)q} \right)-k_{1i}log\left( 1-\rho_{1} \right) \right]$$

In other words, to achieve a particular posterior $q'$, given $\rho_{1}$, $\rho_{2}$ and $q$, the counts $k_{1}$ and $k_{2}$ trade off with each other in a *linear* fashion.

This result gives us a valuable new operation: we can re-express one channel’s evidence count in terms of the other’s, creating a *composite evidence score*. In practice, this converts a count of vocalization recordings into the equivalent number of database entries (or vice versa), which is useful to understand how the hidden mimicry probabilities for each species relates to research attention (Figure S1).

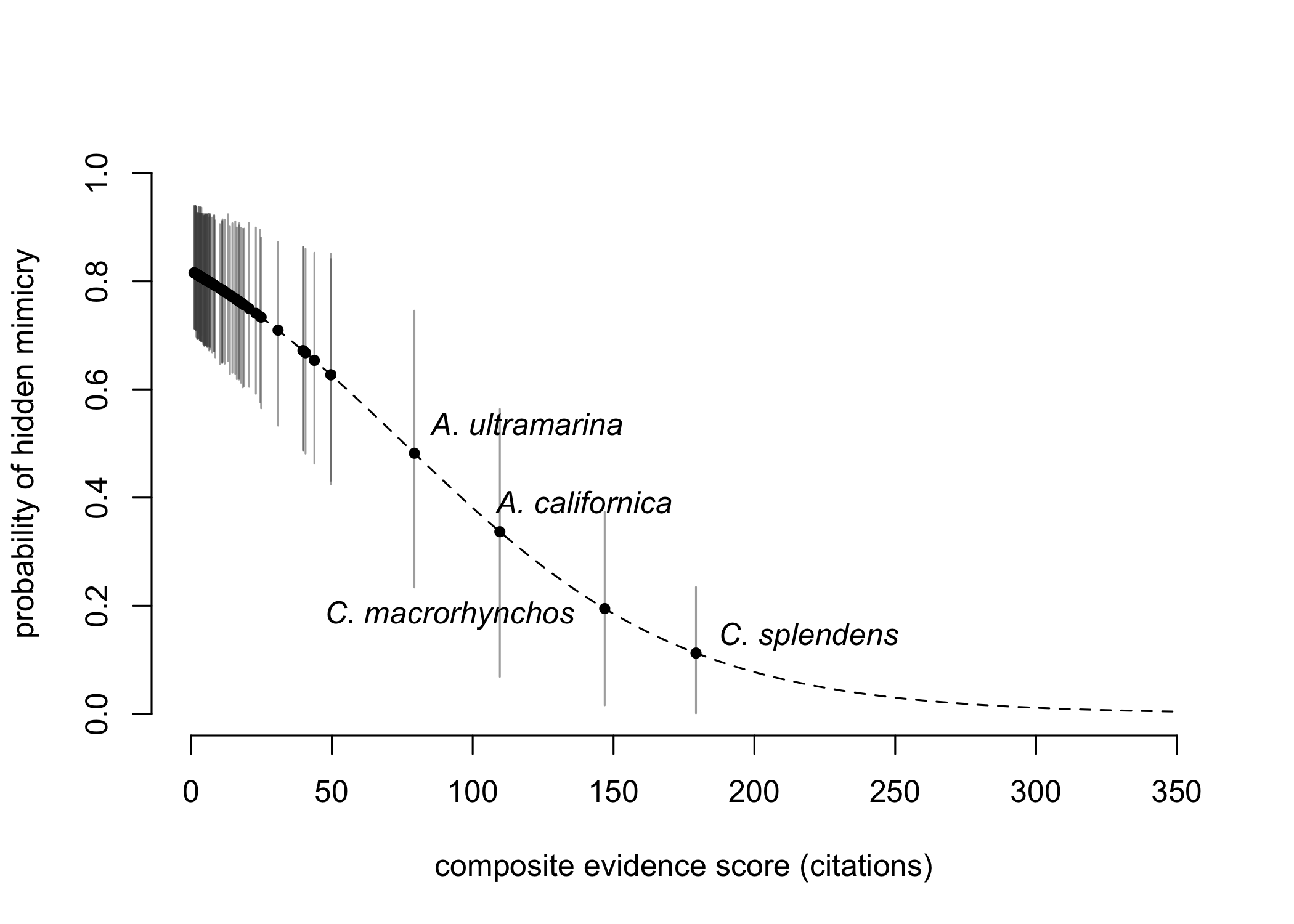

Figure S1. Species-specific posterior predictions from the two-source model (means, with 89% HPDI) plotted against the composite evidence score (as measured in citation equivalents). The dotted line indicates the analytical relationship between these variables.

### Predicting hidden mimicry for specific species

Using the combined model, we can calculate posterior probabilities of hidden mimicry for each species individually. So updated, the model predicts which birds are *least* likely to be hidden mimics (Table S4).

Table S4: Species-specific counts of secondary and primary source evidence and posterior probabilities of hidden mimicry from the two-source model, showing the ten least likely hidden mimics.

| English_name | corvid_database_entries | primary_count | pr_mimic |
| --- | --- | --- | --- |
| House Crow | 188 | 180 | 0.112 (0.105) |
| Large-billed Crow | 133 | 343 | 0.195 (0.135) |
| California Scrub-Jay | 105 | 148 | 0.337 (0.162) |
| Transvolcanic Jay | 81 | 19 | 0.482 (0.160) |
| Azure-winged Magpie | 43 | 88 | 0.627 (0.130) |
| Unicolored Jay | 47 | 39 | 0.627 (0.131) |
| Yellow-billed Chough | 30 | 171 | 0.654 (0.122) |
| Pied Crow | 29 | 143 | 0.668 (0.118) |
| Fish Crow | 29 | 133 | 0.671 (0.118) |
| Mexican Jay | 31 | 107 | 0.672 (0.118) |

Generally speaking, these are birds with large amounts of both primary and secondary evidence, but no reports of mimicry. Conversely, the model nominates these ten birds as most likely to be hidden mimics (Table S5).

Table S5: Species-specific counts of secondary and primary source evidence and posterior probabilities of hidden mimicry from the two-source model, showing the ten most likely hidden mimics. Because of a lack of available evidence, the model returns values very close to the family-wide posterior estimate of q.

| English_name | corvid_database_entries | primary_count | pr_mimic |
| --- | --- | --- | --- |
| Kashmir Nutcracker | 0 | 1 | 0.816 (0.071) |
| Hooded Treepie | 0 | 3 | 0.815 (0.072) |
| Violet Crow | 0 | 6 | 0.815 (0.072) |
| Bougainville Crow | 0 | 7 | 0.814 (0.072) |
| Collared Crow | 0 | 7 | 0.814 (0.072) |
| White-throated Jay | 0 | 7 | 0.814 (0.072) |
| Andaman Treepie | 0 | 8 | 0.814 (0.072) |
| Bismarck Crow | 0 | 10 | 0.814 (0.072) |
| Sumatran Treepie | 0 | 10 | 0.814 (0.072) |
| Purplish-backed Jay | 0 | 11 | 0.813 (0.072) |

The evidence is extremely sparse, so essentially returns the family-wide frequency in every case.

### Ancestral state reconstruction

Our calculations are useful starting places for future fieldwork focusing on vocal mimicry. Based on our empirical data, we calculated ten imputed datasets and used this for imputed ancestral state reconstruction using the posterior estimates from the two-source model (Figure S2).

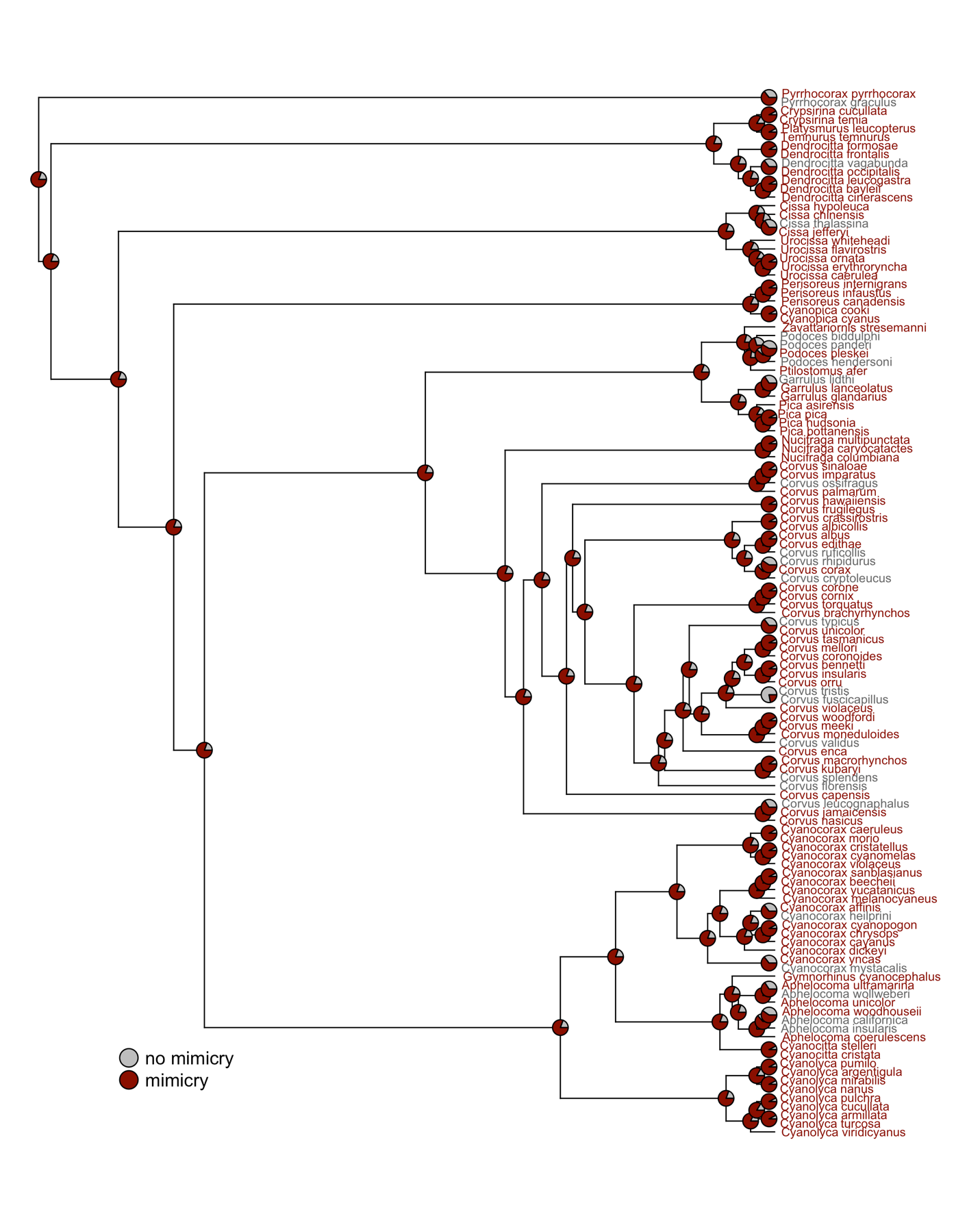

Figure S2. Imputed ancestral state reconstruction using the posterior estimates from the two-source model.

### Robustness Checks

There are a few checks we can make to confirm the validity of the above inferences on the real data.

#### Simulating data from a single and multiple evidence streams

Here’s how a generative simulation would work: for N species, we have a q prior, an average rho, a vector x for their true underlying state (which is what the actual GLM’s would want to study), and a vector D representing the number of recordings for that species. From this, we can compute a pr_obs_ever, which becomes the basis for simulation of Z, the observed mimicry vector.

Supplying the empirical estimates and evidence distributions as simulation input for 1,280 species (x10 the real number of corvids), the posterior estimate of $q$ consistently approaches the true value as each source is added (Figure S3).

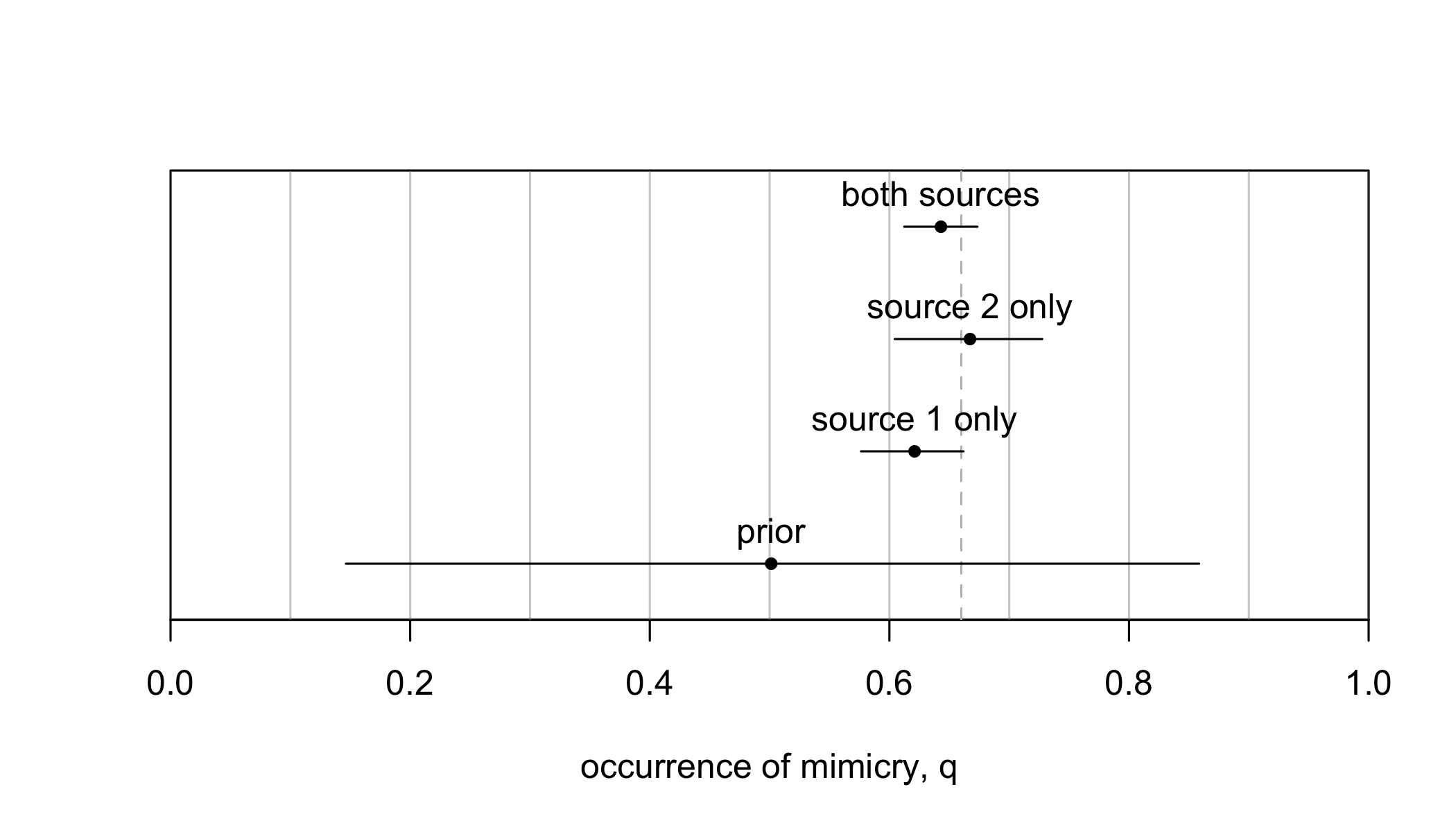

Figures S3: Posterior estimate of the family-wide occurrence of mimicry (means, with 89% HPDI) for a simulation input for ten times the real number of corvid species

#### Filtering low-reliability data

Another potential source of error is our assumption that there are no *false positives* in the detection data. To check the sensitivity of our results to this assumption, Claudia rated the *reliability* of each individual piece of evidence of mimicry, for both primary and secondary sources. For each species showing mimicry, this allows us to calculate the *highest* reliability rating for their phenotype:

| source type | low | medium | high |
| --- | --- | --- | --- |
| xeno-canto | 4 | 2 | 12 |
| secondary | 0 | 4 | 18 |

To check the sensitivity to false positives, we re-analyze the above two-channel model using this information, first under the (extreme) assumption that all “low” reliability records are false (1 -> 0), then under the (very extreme) assumption that *only* “high” reliability records are true.

As the figure below shows, doing so increases the uncertainty around the estimate of occurrence, $q$, but does not substantially shift the evidence away from that rate being high, with a mean occurrence estimate still above 70% in the most conservative false-positive model (Figure S4).

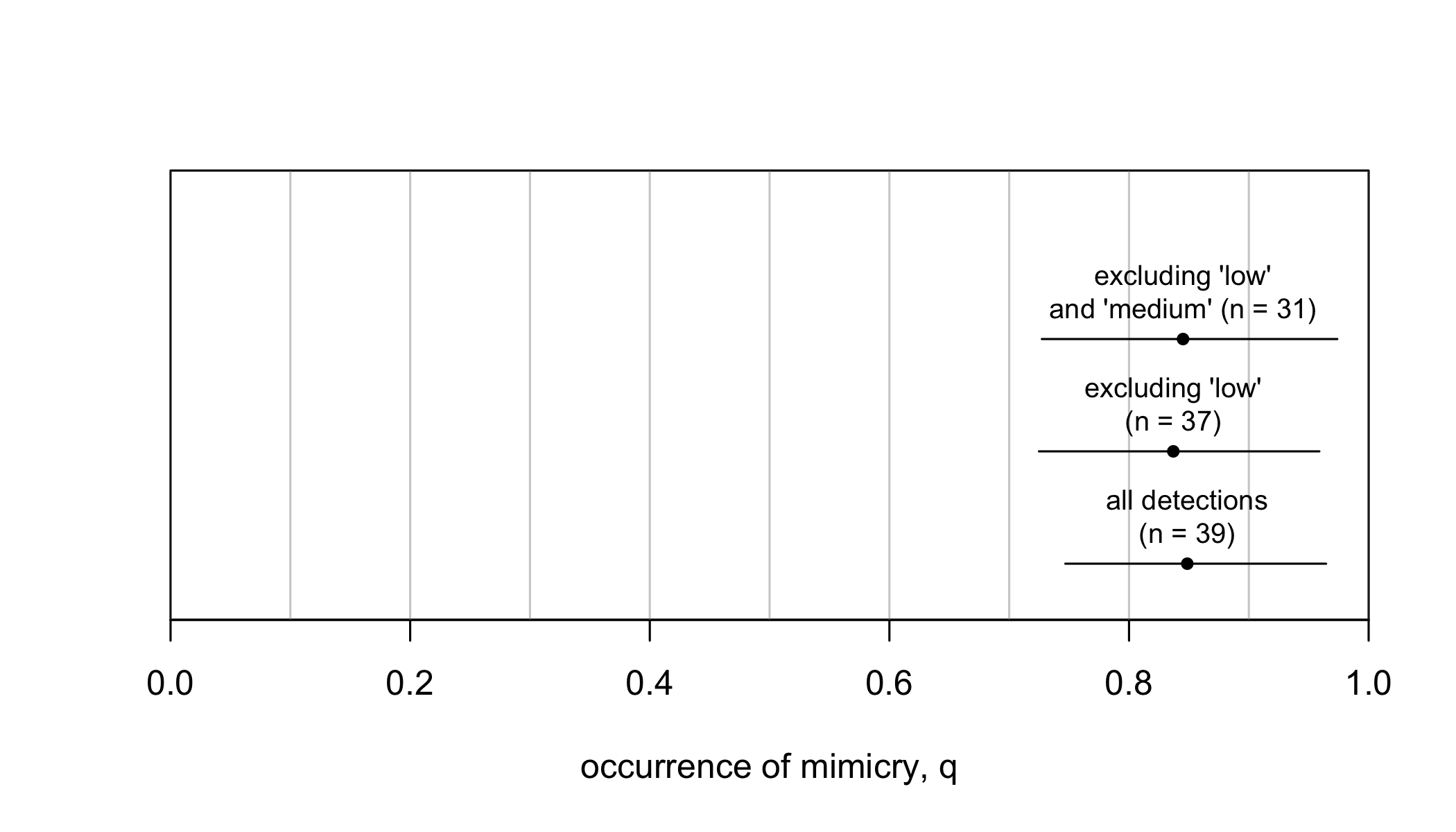

Figure S4. Posterior estimate of the family-wide occurrence of mimicry (means, with 89% HPDI) for different subsets of the data, stratifying by low, medium or high source quality.

#### Different priors on q

A third possible source of sensitivity is the prior assumption on $q$ itself, which in the main text is Beta(2, 2), corresponding to a 50% occurrence rate (mu = 0.5). With only 22 of 128 species known to be mimics in the secondary literature, we might also consider the model results if this prior was also lower.

Setting the prior on q to instead to 20% (Beta(X,Y)), 30%, or 70% produces Figure X. As one might expect, a lower prior on $q$ causes the posterior average to also be lower, but given the weight of the evidence, the posterior estimate of $q$ is still very high, 70% or above (Figure S5). This indicates that the estimated high occurrence rate of mimicry among corvids is not a consequence of prior parameter assumptions.

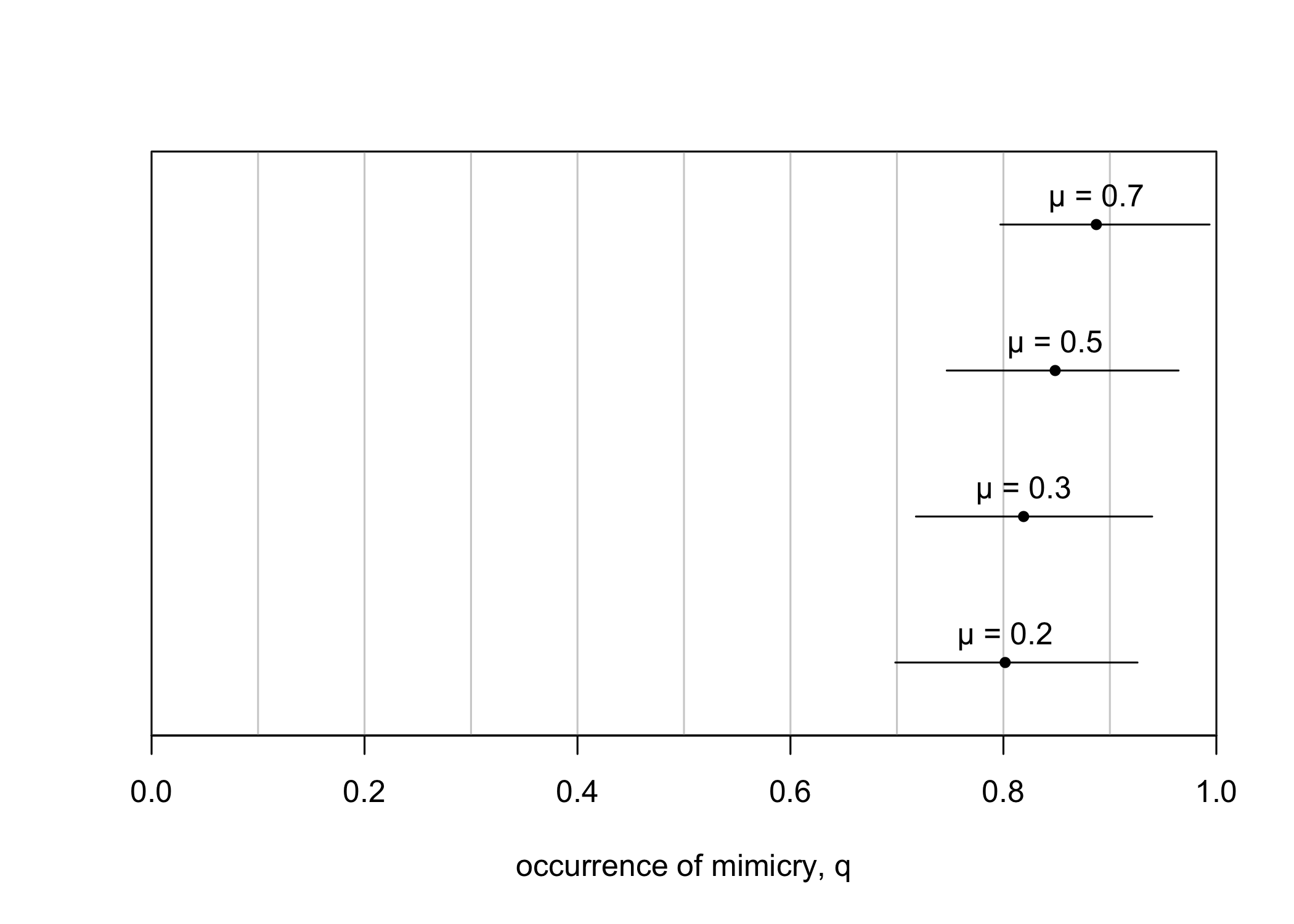

Figure S5. Sensitivity analysis showing four posterior estimates of the occurrence of mimicry (means, with 89% HPDI) with different priors on q, varying from a Beta distribution with an expected value of 0.2 to an expected value of 0.7. In all cases, Beta shape parameters a and b add to 4.

#### Species-specific rho

Different species exhibit mimicry at different rates, within both the primary source recordings and in the secondary literature. We can assess the potential impact on family-wide estimates for q presented in the main text through simulation. This was done in four steps: (1) Observed variation in reporting rates was used to calibrate random simulated values of rho for each of 128 species. Specifically, rates were drawn repeatedly from a Beta(1, 9) distribution. (2) These species rates were then repeatedly re-generated and paired with simulated species-specific mimicry outcomes, drawn randomly from a simulated q set over a grid space between q = 0 and q = 1 at intervals of 0.05. (3) These species-specific parameters (the reporting rates, rho, and the true mimicry status) were used to simulate the main outcome variable, whether mimicry was detected or not detected in a species, for a given number of observations. (4) Finally, we conducted the same statistical analysis on these simulated datasets as described in supplementary section 1, and compared the estimated rate of mimicry occurrence, q, against the true rate for each simulation parameter configuration.

The results of this parameter sweep (Figure S6) indicate there is no systematic bias in the estimation of q for the true sample size of 128 species, though the resolution of these estimates depends on the baseline rate of occurrence itself. As a result, we can conclude that the estimates of q in-text are likely unaffected by variability in detection rates, source quality, or baseline rates of mimicry within specific species.

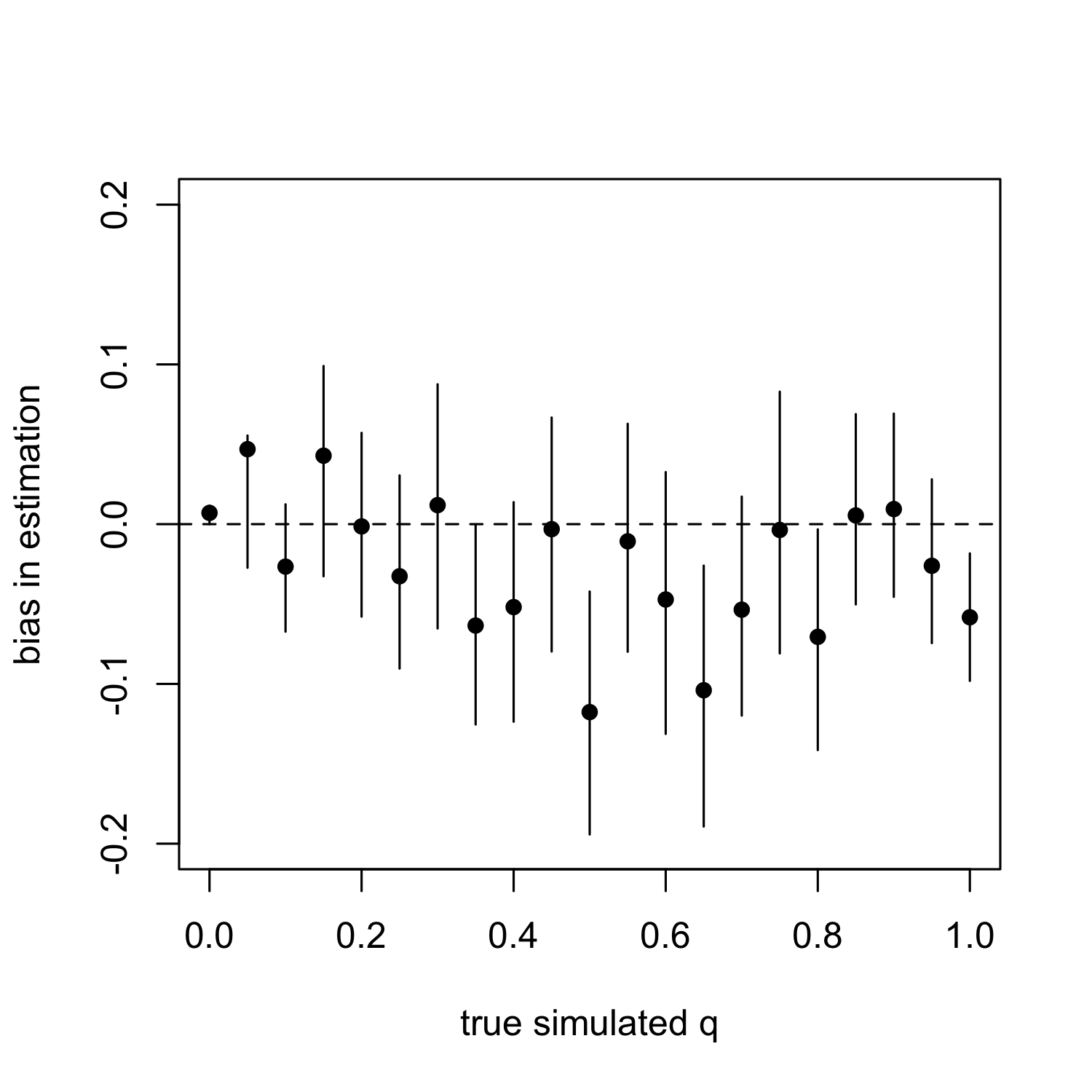

Figure S6. Bias in estimation of family-wide occurrence of mimicry, q, for 21 simulated datasets set at different initial (true) values of q. All bias distributions show means (points) and 89% HPDI. The bias is calculated on the probability scale, subtracting the posterior estimate in q using the single-source MCMC estimates from the true simulated q.
